# Amide I kurtosis maps cardiac amyloid in human tissue by ultrabroadband multiplex coherent anti-Stokes Raman scattering (CARS) microscopy

**DOI:** 10.64898/2026.10.07.757266

**Authors:** Koki Ota, Ting-Hao Chen, Takanori Yamaguchi, Toyokazu Otsubo, Yusuke Murakami, Minori Masaki, Philippe Leproux, Ryo Suzuki, Koichi Node, Hideaki Kano

**Affiliations:** Department of Biosciences and Informatics, Faculty of Science and Technology, Keio University, 3- 14-1 Hiyoshi, Kohoku-ku, Yokohama 223-8522, Japan; Faculty of Medicine, Saga University, 5-1-1 Nabeshima, Saga 849-8501, Japan; Faculty of Science, Kyushu University, 744 Motooka, Nishi-ku, Fukuoka, 819-0395, Japan; International Institute for Integrative Sleep Medicine, University of Tsukuba, 1-1-1 Tennoudai, Tsukuba, Ibaraki, 305-8575, Japan; XLIM Research Institute, CNRS UMR 7252, University of Limoges, 123 Avenue Albert Thomas, F-87060 Limoges Cedex, France

**Keywords:** cardiac amyloidosis, coherent anti-Stokes Raman scattering, CARS, amide I, kurtosis, label-free imaging, second-harmonic generation

## Abstract

Cardiac amyloidosis is caused by deposits of misfolded proteins in the heart and can lead to heart failure. Histological confirmation typically involves staining tissue sections with Congo red and examining them under polarized light for apple-green birefringence; however, achieving an interpretable, label-free, and observer-independent assessment remains challenging. Here we show that Pearson kurtosis of channel intensities within the amide I window provides an interpretable, fitting- free contrast for cardiac amyloid in unstained human cardiac tissue sections imaged by ultrabroadband multiplex coherent anti-Stokes Raman scattering (CARS). In two participants, high- kurtosis regions corresponded with Congo red-positive tissue in adjacent stained sections (descriptive spatial AUCs, 0.79 and 0.73). Across 15 fields from nine participants, including one of these two, kurtosis values did not overlap between severe-deposition and amyloid-negative tissue. Among the descriptors examined, kurtosis showed the largest standardized participant-level separation.

Exploratory multivariate curve resolution yielded a deposit-associated component with aromatic- residue features shared with reagent transthyretin. Second harmonic generation (SHG), acquired simultaneously with CARS, showed a broadly inverse spatial pattern relative to kurtosis, providing complementary architectural contrast. These findings support a label-free approach to detecting cardiac amyloid deposits that combines broadband CARS-derived amide I kurtosis with SHG.

## Introduction

Cardiac amyloidosis is an increasingly recognized cause of heart failure and conduction disease^1,2^. It results from extracellular deposition of amyloid fibrils that impair myocardial compliance, conduction, and tissue architecture^1,2^. Transthyretin amyloidosis (ATTR), in which misfolded transthyretin forms cardiac deposits, is a major cause, particularly in older adults^1–3^. Immunoglobulin light-chain amyloidosis (AL) is another clinically important form^1,2,4,5^. Although their fibrils share a cross-β architecture, they arise from distinct precursor proteins^6–8^.

The availability of disease-modifying therapies has increased the importance of timely diagnosis and accurate amyloid typing^9–11^. Diagnostic evaluation combines echocardiography, cardiovascular magnetic resonance, bone-avid radionuclide imaging, and monoclonal-protein testing^1,2,12^. In selected patients, bone-avid radionuclide imaging together with negative monoclonal-protein testing can establish ATTR cardiac amyloidosis without biopsy^12^. Tissue examination nevertheless remains important when non-invasive findings are inconclusive, a monoclonal gammopathy is present, or the amyloid type remains uncertain. Recent studies using atrial biopsy have demonstrated that amyloid deposition can be detected even in patients without clinically recognized cardiac amyloidosis, including cases with deposits confined to the atria^13,14^. Furthermore, atrial amyloid deposition has been associated with impaired atrial function and heart failure in patients undergoing catheter ablation for atrial fibrillation^15^, highlighting the clinical relevance of assessing amyloid burden.

In tissue available for examination, Congo red staining and apple-green birefringence under polarized light establish amyloid deposition in tissue^16,17^. Identifying the precursor protein requires a separate assessment, usually by immunohistochemistry and, when inconclusive, mass spectrometry^18^. Quantitative molecular imaging of unstained sections could complement histopathology when tissue is already available for examination.

Vibrational spectroscopy offers one route to such analysis. The amide I band, dominated by peptide-backbone carbonyl stretching, is sensitive to protein secondary structure and has long been used to study amyloid formation^19–22^. Previous Raman studies have distinguished amyloid deposits from surrounding human tissue using amide I and fingerprint-region features^23^, but weak spontaneous Raman signals constrain imaging speed. Coherent Raman techniques, including coherent anti-Stokes Raman scattering (CARS), enable faster chemical imaging^24,25^. At each pixel, ultrabroadband multiplex CARS acquires a spectrum spanning the fingerprint region and the CH/OH-stretching regions, from which a Raman-equivalent response can be retrieved^26–30^. This makes it possible to map spatial variation in amide I band shape, but an interpretable, fitting-free contrast for cardiac amyloid has not been established.

Here we show that Pearson kurtosis of channel intensities within the amide I window, a descriptor sensitive to how sharply peaked the spectral profile is, provides an interpretable, fitting-free contrast for mapping cardiac amyloid in unstained human cardiac tissue. We examined its spatial correspondence with Congo red-positive deposits and its relationship to amide I band narrowing in two adjacent-section CARS–histology pairs from two participants. We then assessed cross-participant consistency in 15 fields from nine participants, including one of the two examined in the section-pair analysis. Exploratory multivariate curve resolution–alternating least squares (MCR-ALS) and second harmonic generation (SHG) acquired simultaneously with CARS placed this contrast in broader chemical and architectural contexts.

## Results

### Amide I kurtosis maps tissue regions corresponding to Congo red-positive deposits

Ultrabroadband multiplex CARS imaging of an unstained human cardiac section yielded a Raman- equivalent Im[χ⁽³⁾] response spanning approximately 440–3690 cm⁻¹. The CARS image was registered to the adjacent Congo red-stained serial section from the same participant. Integrated amide I intensity over 1600–1720 cm⁻¹ visualized tissue morphology but showed only weak spatial correspondence with registered Congo red-positive regions (Spearman’s ρ = 0.36; area under the receiver operating characteristic curve (AUC) = 0.66). To derive a fitting-free descriptor from the amide I window, we calculated Pearson kurtosis of the baseline-corrected channel intensities within the same interval at each pixel. This metric is invariant to uniform multiplicative intensity scaling and therefore describes the distribution of intensities largely independently of absolute signal magnitude. Compared with integrated intensity, the kurtosis map showed closer spatial correspondence with the Congo red- positive mask registered from the adjacent section (ρ = 0.53; AUC = 0.79; Fig. 1). Analysis of an unstained CARS section from a second participant, who was also included in the nine-participant comparison in Fig. 3, showed similar but weaker correspondence with Congo red-positive tissue in the registered adjacent section (ρ = 0.39; AUC = 0.73; Supplementary Results). Together, the two adjacent-section pairs showed consistent spatial correspondence between high amide I kurtosis and Congo red-positive tissue, prompting us to examine the spectral basis of this contrast.

**Figure 1.**
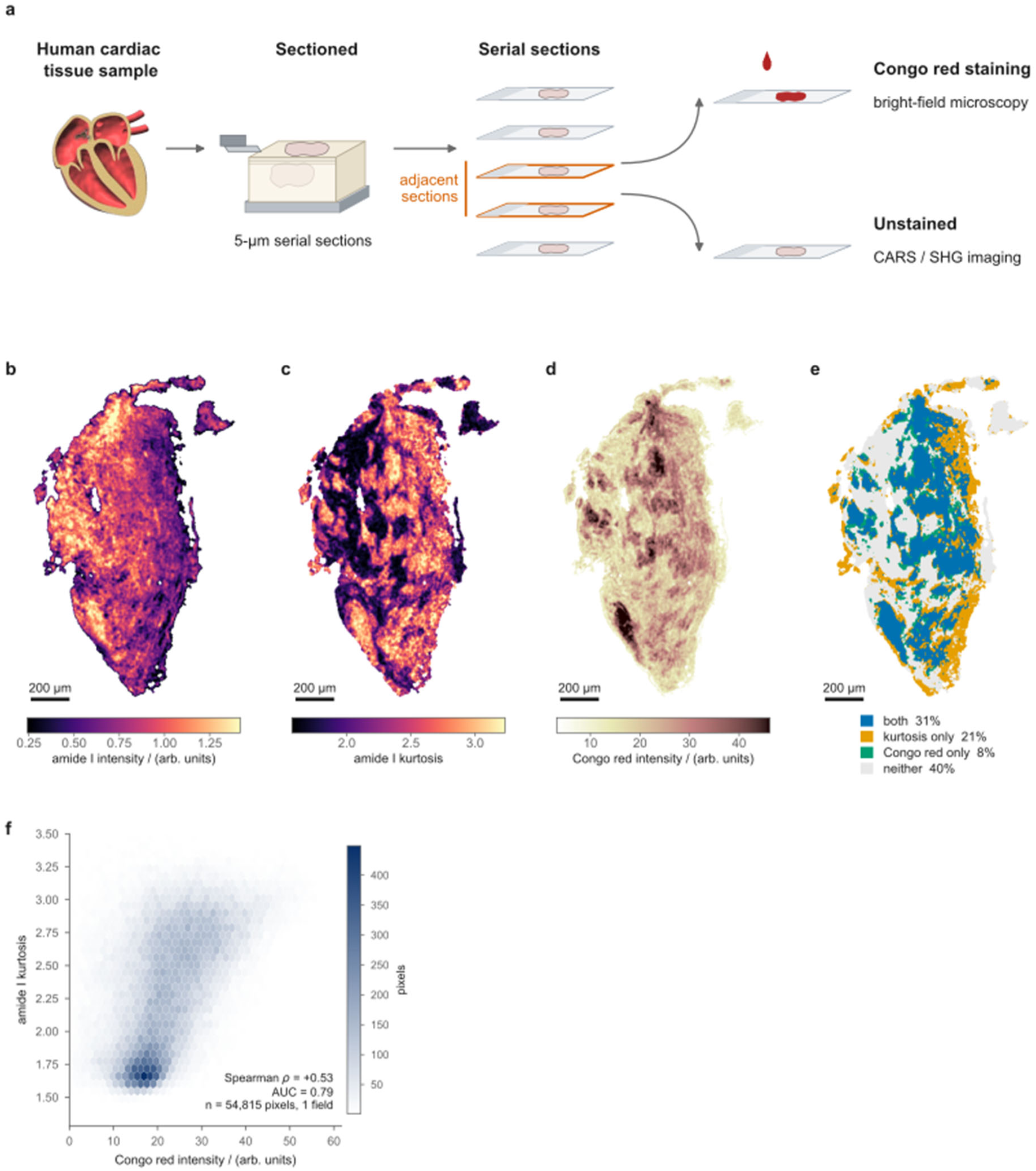
Amide I kurtosis maps tissue regions corresponding to Congo red-positive deposits. (a) Adjacent- section workflow. Serial 5-µm cardiac tissue sections were assigned to Congo red histology and unstained CARS/SHG imaging. (b) Integrated amide I intensity over 1600–1720 cm⁻¹ and (c) Pearson kurtosis of baseline- corrected channel intensities within the same window and CARS field. (d) Bright-field Congo red image from the adjacent stained section after registration to the CARS grid. (e) Overlay of independently thresholded kurtosis and Congo red masks: both positive, 31%; kurtosis only, 21%; Congo red only, 8%; neither, 40%. (f) Joint distribution of paired pixel values (Spearman’s ρ = 0.53; spatial AUC = 0.79); color denotes pixel count. These spatial statistics describe correspondence across adjacent sections (n = 54,815 spatially correlated pixels in one field). Scale bars, 200 µm.

**Figure 2.**
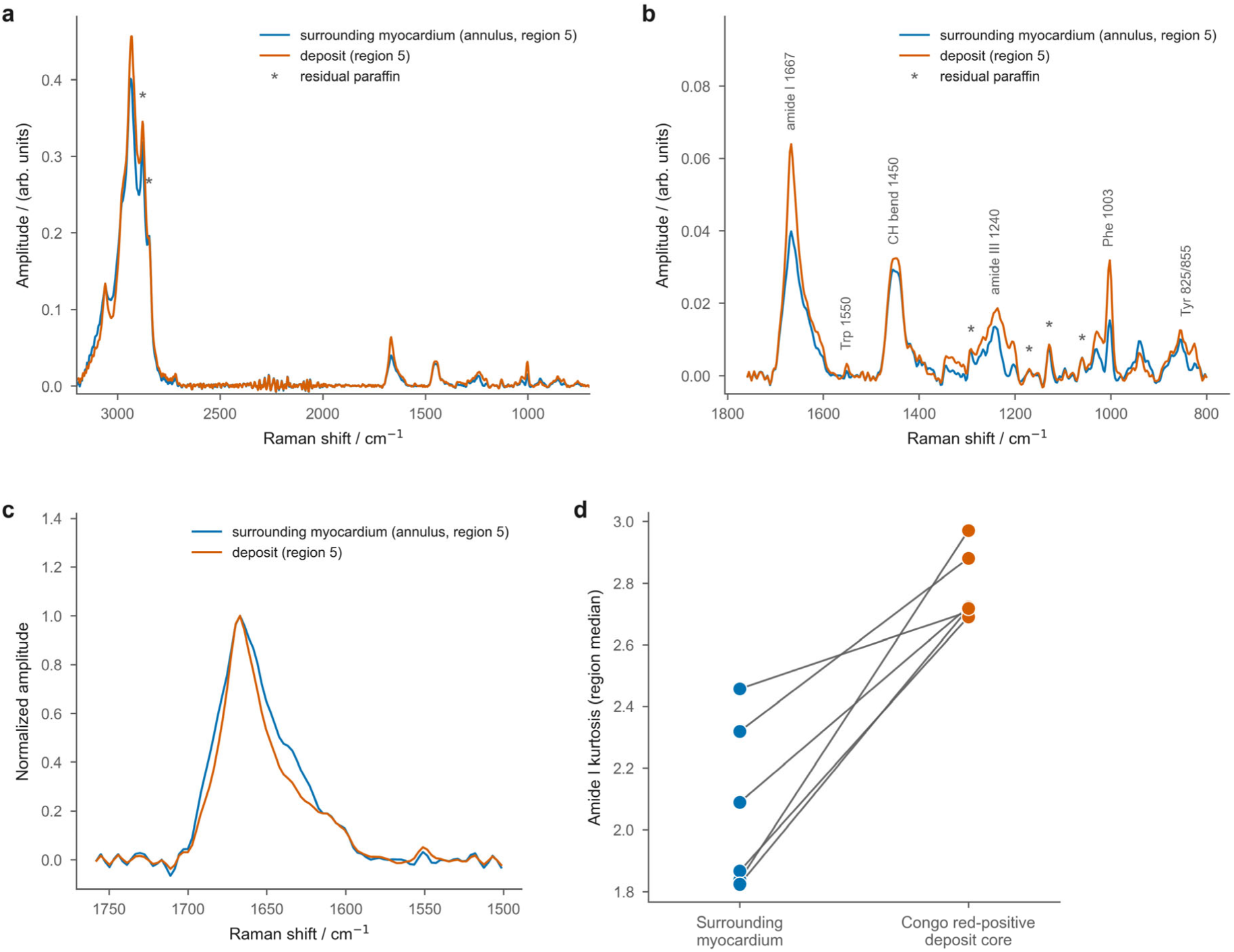
Amide I kurtosis captures deposit-associated changes in band shape. (a–c) Raman-equivalent Im[χ^(3)^] mean spectra from deposit-associated region 5 (orange) and its paired surrounding-myocardium annulus (blue), 80–240 µm outside the registered deposit boundary, shown over (a) the full measured range, (b) the fingerprint and amide I regions and (c) the expanded amide I region after baseline correction and normalization to peak intensity. Asterisks in (a) and (b) mark bands with substantial contributions from residual paraffin. (d) Regional median amide I kurtosis for all six deposit–annulus pairs in the first registered section; lines connect paired regions. All six pairs came from one participant.

**Figure 3.**
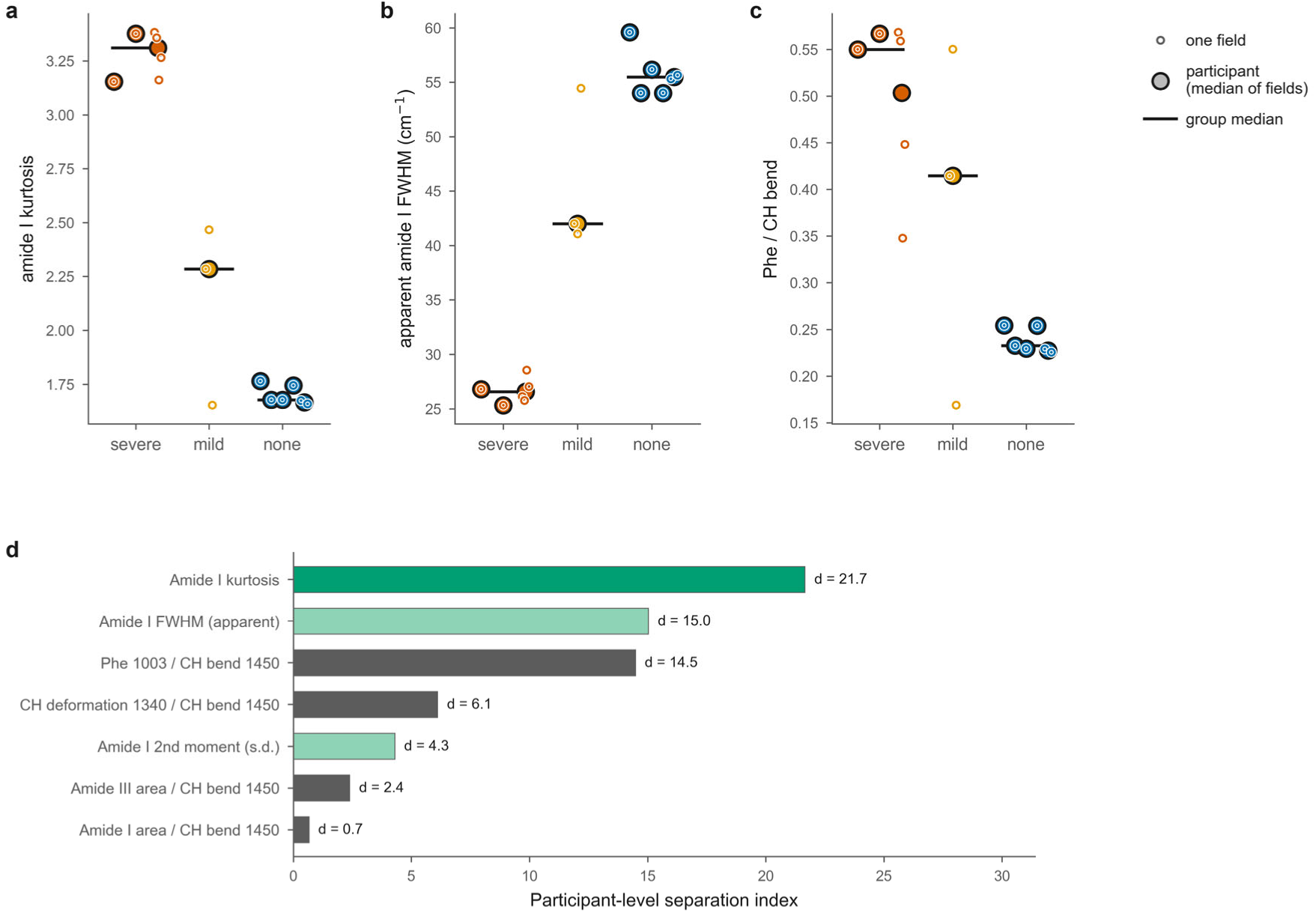
Amide I kurtosis shows the largest standardized participant-level separation among the spectral descriptors examined. Fifteen fields were acquired from nine participants: four with ATTR cardiac amyloidosis (three with severe deposition, six fields; one with mild deposition, three fields) and five amyloid-negative participants without ATTR (six fields). Deposition grades were assigned by histopathological assessment. (a) Amide I channel-intensity kurtosis, (b) apparent amide I FWHM obtained by pseudo-Voigt fitting of each field- mean spectrum and (c) the phenylalanine (1003 cm⁻¹)/CH bending (1450 cm⁻¹) ratio. Small open symbols show field values (medians of pixel values in a and c; fitted values of the field-mean spectrum in b); larger black- outlined symbols show participant medians. Orange, severe; yellow, mild; blue, amyloid-negative. For kurtosis, severe and amyloid-negative participant medians differed (exact two-sided Mann–Whitney U test, P = 0.036). (d) Exploratory separation index for seven spectral descriptors, calculated from participant-level values. For each descriptor, the absolute difference between the severe and amyloid-negative group medians was divided by the pooled within-group standard deviation. The participant with mild deposition was excluded. The ranking is descriptive given the small number of participants.

### Amide I kurtosis captures deposit-associated changes in band shape

To examine the spectral basis of the kurtosis contrast, we compared six regions in the unstained CARS section corresponding to Congo red-positive deposits in the adjacent stained section with paired regions of surrounding myocardium (Fig. 2). As an example, mean spectra from deposit-associated region 5 and its paired surrounding-myocardium annulus are shown across the full measured range, the fingerprint region and the expanded amide I region (Fig. 2a–c). After baseline correction and normalization, the deposit-associated amide I band appeared narrower than that of the paired myocardium (Fig. 2c). Regional median amide I kurtosis was higher in deposit-associated tissue in all six registered region pairs (Fig. 2d). Apparent amide I full width at half maximum (FWHM) was smaller in deposit-associated tissue than in the paired annulus for all six regions from the discovery participant and in five of six from the second participant. In the second participant the median paired decrease was approximately 18 cm⁻¹ (Supplementary Fig. S1). Peak-position analysis did not establish a reproducible shift. Thus, kurtosis provided a fitting-free readout of the deposit-associated amide I band-shape change, supported by FWHM narrowing in both participants.

### Amide I kurtosis separates severe deposition from amyloid-negative tissue across participants

To assess cross-participant consistency, we analyzed 15 fields from nine participants, including one of the two participants in the adjacent-section analysis. Four had ATTR cardiac amyloidosis (three with severe deposition, six fields; one with mild deposition, three fields) and five were amyloid- negative (six fields). Field-level kurtosis values did not overlap between the severe and amyloid- negative groups (3.15–3.38 versus 1.66–1.76; Fig. 3a), and participant-level medians retained this separation (exact two-sided Mann–Whitney U test, P = 0.036). Apparent amide I FWHM was smaller in severe-deposition fields than in amyloid-negative fields (25–29 versus 54–60 cm⁻¹; Fig. 3b), consistent with the deposit-associated narrowing observed in both registered section pairs (Supplementary Fig. S1). The Phe (1003 cm⁻¹)/CH bending (1450 cm⁻¹) ratio also separated the groups (Fig. 3c). Among the descriptors examined, kurtosis showed the largest standardized participant-level separation in this exploratory comparison (Fig. 3d).

The three fields from the participant with mild deposition showed heterogeneous kurtosis values (Fig. 3a). To test whether group separation remained detectable with restricted spectral coverage or poorer signal quality, we varied the amide I analysis window and artificially reduced the signal-to- noise ratio (SNR) of each field. Separation persisted across all tested windows except the narrowest one (Supplementary Fig. S2). Degradation to the lowest native field-level SNR in the cohort (2.54) narrowed the gap between severe and amyloid-negative kurtosis ranges (Supplementary Fig. S3).

Although the mild case and SNR dependence do not support three-grade classification or acquisition-independent performance, kurtosis provided a clear, fitting-free contrast between severe-deposition and amyloid-negative tissue in the data as acquired.

### MCR-ALS resolves a deposit-associated component with transthyretin-consistent features

To examine whether high-kurtosis regions also showed deposit-associated spectral features outside the amide I window, we applied exploratory four-component MCR-ALS^31^. Among the resolved components, we examined the one most closely associated spatially with the amide I kurtosis map (Spearman’s ρ = 0.84). This component also corresponded with registered Congo red-positive tissue (ρ = 0.59; AUC = 0.82; Fig. 4). Its spectrum contained an amide I feature at 1667 cm⁻¹, a phenylalanine-associated band near 1003 cm⁻¹, a tryptophan-associated feature near 1550 cm⁻¹, and a tyrosine-associated doublet near 825 and 855 cm⁻¹, with the 825 cm⁻¹ band being more prominent.

**Figure 4.**
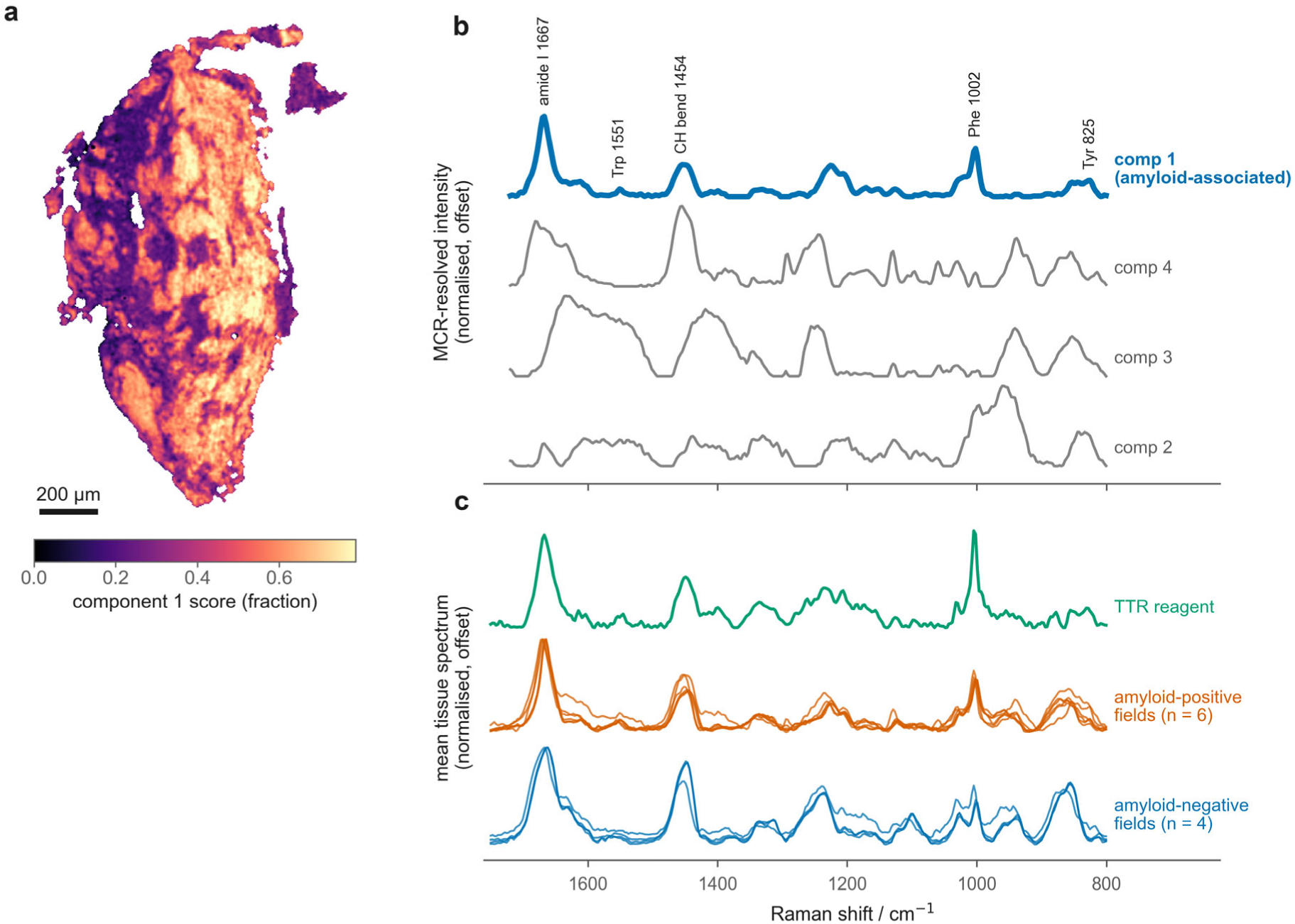
MCR-ALS resolves a deposit-associated component with transthyretin-consistent spectral features. (a) Score map of the component selected for its spatial correspondence with amide I kurtosis. (b) Four resolved component spectra over the fingerprint and amide I regions, normalized and vertically offset. The selected component (blue) has features near 825 and 855, 1003, 1550 and 1667 cm⁻¹. Its score map corresponded with registered Congo red-positive tissue (Spearman’s ρ = 0.59; spatial AUC = 0.82). (c) Qualitative comparison of a separately acquired reagent transthyretin spectrum with individual field-mean tissue spectra. The tissue spectra comprise one discovery field, three fields from participants with severe deposition, two from the participant with mild deposition and four fields from three amyloid-negative participants. Before each field mean was calculated, the tissue data were numerically degraded to the lowest native SNR among these ten fields and spatially smoothed (σ = 1 pixel). The shared fingerprint-region features support compositional compatibility with TTR- containing deposits but do not establish precursor identity or amyloid type. Scale bar, 200 µm.

Several of these fingerprint-region features were also present in the separately acquired reagent transthyretin spectrum and in field-mean tissue spectra from the discovery participant and the additional cohort (Fig. 4c). Together, deposit localization, amide I narrowing and the fingerprint resemblance to reagent transthyretin are consistent with a contribution from TTR-containing deposits. The MCR-ALS comparison in the second registered section is summarized in the Supplementary Results, while targeted aromatic-region maps from the discovery field are shown in Supplementary Fig. S4. These features are shared by other proteins, and the spatial maps were derived from the same CARS dataset, so they do not independently establish precursor identity or amyloid type.

Nevertheless, the TTR-consistent fingerprint pattern extends the kurtosis finding beyond amide I band shape and supports its association with ATTR deposits.

### Simultaneous SHG reveals complementary collagen-associated architectural contrast

Fibrillar collagen is a strong source of SHG in myocardium,^32,33^ and we interpreted the signal in this field as predominantly collagen-derived. SHG and CARS images were acquired simultaneously from the same unstained section and field shown in Fig. 1 (Fig. 5a,b). The SHG intensity map showed a broadly inverse spatial pattern relative to the amide I kurtosis map (Spearman’s ρ = −0.68). SHG intensity was also lower within registered Congo red-positive regions than outside them. Together, these measurements show that amyloid-associated changes in amide I band shape coincide with reduced SHG from collagen-rich myocardial structures in this field.

**Figure 5.**
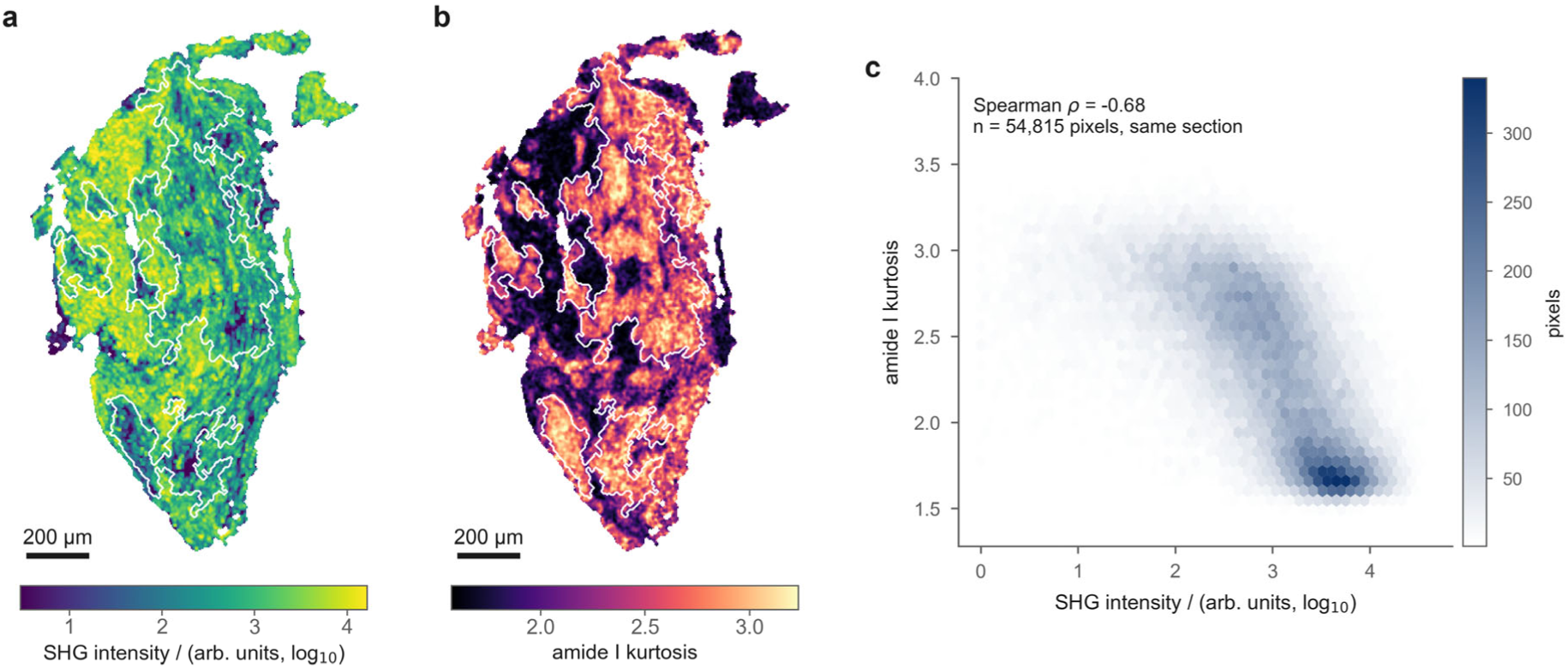
Simultaneously acquired SHG provides complementary architectural contrast. (a) SHG intensity, displayed on a log₁₀ scale, and (b) amide I kurtosis acquired from the same unstained section and field. Identical white outlines in (a) and (b) delineate Congo red-positive regions registered from the adjacent stained section (Otsu threshold; connected regions ≥ 400 pixels). (c) Joint distribution of pixel-wise kurtosis and log₁₀- transformed SHG intensity (Spearman’s ρ = −0.68; n = 54,815 spatially correlated pixels); color denotes pixel count. Median raw SHG intensity was 589 arb. units within registered regions and 1,539 arb. units outside them. Scale bars, 200 µm.

## Discussion

This study identifies amide I kurtosis as an interpretable, fitting-free CARS contrast associated with cardiac amyloid deposits in unstained tissue sections. In two participants with registered histology, high-kurtosis regions corresponded spatially with Congo red-positive deposits identified in adjacent sections. In the nine-participant field analysis, kurtosis values did not overlap between severe-deposition and amyloid-negative fields and showed the largest standardized participant-level separation among the descriptors examined. Higher kurtosis accompanied a smaller apparent amide I FWHM, whereas no reproducible peak shift was resolved. Integrated amide I intensity showed weaker spatial correspondence with Congo red-positive tissue. Together, these findings support kurtosis as a fitting-free readout of deposit-associated changes in amide I band shape.

Kurtosis summarizes the distribution of intensity values within a fixed spectral window without explicitly accounting for their positions along the wavenumber axis. It is therefore not a direct measure of linewidth, β-sheet content or a specific secondary structure. Within the predominantly single-peaked amide I envelopes observed here, however, higher kurtosis accompanied a smaller apparent FWHM. A more homogeneous distribution of peptide carbonyl environments in the ordered cross-β structure of amyloid is one possible explanation, consistent with previous vibrational studies^19,22^. Biochemical composition, local environment and overlapping bands may also affect the observed shape. The severe–amyloid-negative separation also weakened after numerical reduction of the signal-to-noise ratio. Kurtosis is thus a useful fitting-free descriptor under the acquisition and analysis conditions examined, but its structural interpretation and performance require further testing.

MCR-ALS placed the kurtosis contrast in a broader chemical context. The component selected for its spatial correspondence with kurtosis also corresponded with registered Congo red-positive tissue. Its spectrum contained an amide I feature at 1667 cm⁻¹ and aromatic-residue-associated features near 1003, 1550, 825 and 855 cm⁻¹. The lower-wavenumber member of the tyrosine-associated doublet was more prominent. The doublet arises from Fermi resonance, and its relative intensities are sensitive to the environment of the phenolic hydroxyl group^34^. In amorphous aggregates of the FUS low-complexity domain, a lower conventional I₈₅₀/I₈₃₀ ratio likewise accompanied amide I band narrowing^35^. This qualitative parallel supports the interpretation that changes in tyrosine side-chain and peptide-backbone environments occur together in the amyloid-associated tissue examined here.

Several fingerprint-region features were also present in a separately acquired reagent transthyretin spectrum. Together with deposit localization and amide I narrowing, this resemblance supports a TTR contribution to the deposit-associated CARS signal in the ATTR tissue examined here.

In the cardiac section examined here, we interpret SHG as arising predominantly from fibrillar collagen, a well-established source of SHG in myocardium^32,33^. Cardiac myosin can also contribute,^36^ and SHG intensity depends on fiber orientation, organization, packing and optical phase matching as well as collagen abundance. The images therefore do not quantify collagen content. Nevertheless, the broadly inverse SHG–kurtosis relationship and lower SHG intensity in registered Congo red-positive regions support a local alteration of collagen-rich myocardial architecture accompanying amyloid deposition. SHG thus places the CARS-derived spectral contrast within its structural context.

CARS imaging does not require removal of tissue from the measured section, unlike laser microdissection–mass spectrometry, which samples deposit-containing material for amyloid typing. The same section could then be stained, allowing direct comparison of CARS maps with histology when biopsy material is limited. With further development, a fiber-based CARS probe^37^ could bring molecular assessment to the endocardial surface during biopsy and help guide tissue sampling.

These findings remain preliminary. Histology-registered analyses were limited to two participants, each with one pair of adjacent sections; section-to-section variation and registration uncertainty therefore constrain pixel-level comparisons. The additional cohort was small and, except for one participant, lacked field-matched histology, and the participant with mild deposition showed heterogeneous results. Field selection, anatomical site, tissue processing, and acquisition session may also have influenced the comparisons. Same-section CARS followed by staining, standardized acquisition and quality control, predefined thresholds and blinded validation in an independent participant-level cohort are needed to establish generalizability and diagnostic performance. Within these limits, amide I kurtosis provides an interpretable, fitting-free readout of amyloid-associated band shape in *ex vivo* cardiac tissue, complemented by the chemical and architectural context from broadband spectral decomposition and simultaneous SHG.

## Methods

### Human cardiac tissue samples and ethical approval

Human right atrial tissue from ten participants was analyzed. One discovery participant and one of the nine additional participants each contributed one unstained section registered to an adjacent Congo red-stained section. Six deposit-associated regions meeting the area criterion were analyzed in each section pair. The discovery fields were selected with reference to the corresponding histology and were not selected blind. Nine additional participants contributed 15 CARS fields. Four had ATTR cardiac amyloidosis, and five were amyloid-negative participants without ATTR. Pathological assessment of Congo red-stained sections classified three participants as having severe deposition (six fields), one as having mild deposition (three fields) and five as amyloid-negative (six fields).

The study was approved by the Clinical Research Ethics Committee of Saga University Hospital (approval nos. 2020-01-01 and 2020-09-01). Research at the Faculty of Science and Technology and Graduate School of Science and Technology, Keio University was conducted under institutional permission nos. 2026-047 and 2026-048. Information about the present analysis was publicly disclosed, and participants were given the opportunity to opt out. The requirement for written informed consent was waived by the Clinical Research Ethics Committee of Saga University Hospital owing to the retrospective nature of the study. All procedures were conducted in accordance with the Declaration of Helsinki and applicable institutional and national regulations. Samples and images were de-identified before analysis.

### Tissue processing and serial sectioning

Tissue was fixed in paraformaldehyde, embedded in paraffin and cut into 5-µm-thick serial sections. Adjacent sections were used for Congo red histology or label-free CARS and SHG imaging. Sections used for optical imaging were deparaffinized and examined without staining.

### Congo red staining and conventional microscopy

Paraffin sections were heated at 55 °C for 15 min, immersed three times in xylene for 15 min each and then immersed three times in 70% ethanol for 10 min each before rehydration in running water. Sections were stained with hematoxylin for 5 min and Congo red for 10 min, washed and briefly immersed in lithium carbonate solution. Amyloid status was determined by pathological assessment of Congo red staining, including apple-green birefringence under polarized light^16,17^. Bright-field images of the Congo red-stained sections used for registration were acquired using a conventional inverted optical microscope.

### Ultrabroadband multiplex CARS and SHG microscopy

Measurements were performed using a custom multimodal nonlinear optical microscope based on previously described systems^27,38,39^. A dual-fiber synchronized source (OPERA HP2, Leukos, Limoges, France) delivered 1064-nm master pulses (approximately 50 ps, 5 MHz) and broadband near-infrared supercontinuum Stokes pulses. The two pulses were overlapped spatially and temporally, introduced coaxially into an inverted microscope and focused with an objective of numerical aperture 1.27. Powers at the specimen were 80 and 50 mW for the pump and Stokes beams, respectively. The transmitted CARS signal was collected with an objective of numerical aperture 0.6, dispersed by a spectrograph (LS-785, Princeton Instruments, Trenton, NJ, USA) and detected with a CCD camera (BLAZE 100HR, Princeton Instruments) controlled by LightField software. Samples were scanned with an LP-200 piezoelectric stage (Mad City Labs, Madison, WI, USA) mounted on a motorized stage. The discovery field used 4-µm spatial sampling over a 1.8 mm × 1.8 mm field; additional fields measured 100 µm × 100 µm. SHG was excited by the pump beam and dispersed by a spectrograph (SpectraPro 300i, Princeton Instruments, Trenton, NJ, USA) and detected with a CCD camera (PIXIS- B, Princeton Instruments). CARS and SHG data in Fig. 5 were acquired simultaneously from the same section and field. The exposure time for each pixel was typically 20 ms. The Im[χ^(3)^] response was retrieved from the CARS spectra by the maximum entropy method^40,41^. Raman shift was calibrated against polystyrene beads (Polysciences).

### Image registration between CARS and histology

The Congo red image was registered to the CARS grid using corresponding landmarks and a thin-plate-spline transform in Fiji/ImageJ^42^ with BigWarp^43.^ The same workflow was applied to the second section pair.

### Amide I intensity and kurtosis analysis

The amide I interval was defined as 1600–1720 cm^−1^. A straight baseline connecting the interval endpoints was subtracted at each pixel, and integrated amide I intensity was calculated. Tissue pixels were identified from CH-stretch intensity, and the same tissue mask was used for all amide I analyses.

For each retained pixel, Pearson kurtosis K was calculated from N baseline-corrected channel intensities Iᵢ within the amide I interval:

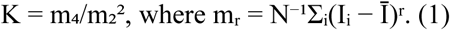

No finite-sample bias correction or subtraction of 3 was applied. Because K is calculated from intensity values without weighting by wavenumber, it is invariant to channel order and uniform multiplicative scaling and differs from intensity-weighted spectral kurtosis along the wavenumber axis. It describes the distribution of intensity values within the selected interval rather than spectral width directly.

### Amide I width and sensitivity analyses

For each registered dataset, Congo red-positive regions with an area of at least 6,400 µm² were analyzed separately. Each deposit-associated region was paired with a surrounding-myocardium annulus 80–240 µm from its boundary. Mean spectra were calculated for each region and its paired annulus. To estimate apparent amide I FWHM, a straight line joining the endpoints of the 1580–1750 cm⁻¹ window was first subtracted from each mean spectrum. A pseudo-Voigt profile with a free residual linear baseline was then fitted by least squares, and FWHM was determined from the fitted profile. Fits whose width reached either parameter bound (5.5 or 169 cm⁻¹) were excluded. The width is termed apparent because it was not deconvolved for instrumental resolution. The same procedure was applied to field-mean spectra from the additional cohort. The intensity-weighted spectral standard deviation was calculated from the second central moment within 1600–1720 cm⁻¹. Kurtosis window sensitivity was assessed using intervals centered at 1660 cm⁻¹ with half-widths of 30–100 cm⁻¹.

### Between-participant and SNR sensitivity analyses

Pixel-wise spectral descriptors were summarized by the median over tissue pixels within each additional field. Apparent amide I FWHM was instead obtained by fitting the mean tissue spectrum of each field, as described above. For participants contributing more than one field, the participant-level value for each descriptor was the median of its field-level values. Native SNR was estimated using the same procedure for all fields. For the sensitivity analysis, independent simulated noise was added to each spectrum until the field-level SNR reached 2.54, the lowest native SNR observed among the 15 additional-cohort fields. The spectral metrics were then recalculated. This exploratory analysis was not used to define a diagnostic threshold.

### Congo red spatial correspondence

Registered Congo red intensity and amide I kurtosis values were paired within the common tissue mask. Otsu thresholding was applied independently to the two images to generate the concordance overlay in Fig. 1e. The spatial AUC was calculated using the binary Congo red mask as the reference label. Spearman correlations and pixel-level AUCs were treated as descriptive spatial- correspondence measures because neighboring pixels were not independent.

### MCR-ALS analysis

For multivariate analysis, the spectral region spanning the fingerprint and amide I bands shown in Fig. 4b was unfolded into a matrix X with pixels as rows and Raman shifts as columns. MCR-ALS decomposed the data according to

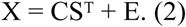

Here C contains component score maps, S contains resolved spectra and E is the residual matrix. A four-component model was initialized from absolute principal-component loadings, with non-negativity constraints on scores and spectra. The deposit-associated component was selected by its spatial correspondence with kurtosis. Because the MCR-ALS and kurtosis maps were derived from the same CARS dataset, their mutual correspondence was treated as internal consistency; comparison with registered Congo red provided spatial context but not independent molecular identification.

### Aromatic-residue-associated maps and reagent transthyretin

After local linear baseline correction, the phenylalanine-associated band over 996–1010 cm⁻¹ and the tryptophan-associated region over 1540–1565 cm⁻¹ were integrated. Each band area was divided by the CH bending-band area near 1450 cm⁻¹ to generate the maps in Supplementary Fig. S4a,b. The tyrosine doublet was calculated as I_855_/(I_825_ + I_855_) using 818–832 and 848–862 cm⁻¹ windows. For the additional-cohort descriptors (Fig. 3), band areas at 996–1010 (phenylalanine), 1225–1265 (amide III), 1320–1360 (CH deformation), 1440–1462 (CH bending) and 1600–1720 cm⁻¹ (amide I) were each integrated after a local linear baseline between flanking windows, and the phenylalanine, amide I, CH- deformation and amide III areas were divided by the CH bending-band area. A separately acquired CARS reference spectrum of reagent transthyretin was phase retrieved, baseline-corrected with a minimum-filter spline after Savitzky–Golay smoothing and normalized to its maximum within 800– 1750 cm⁻¹ before qualitative comparison with the deposit-associated MCR-ALS component and field- mean tissue spectra. No classifier or prespecified similarity threshold was applied.

### SHG image analysis

SHG images were reconstructed by fitting a Gaussian function to the SHG spectrum at each pixel and smoothed with a 3 × 3 binomial kernel. SHG intensities below 1 arb. unit were set to 1 arb. unit before log₁₀ transformation. Spearman’s correlation with amide I kurtosis was calculated from the transformed intensities within the common tissue mask. The log₁₀ transformation preserved the rank order above the clipping threshold; clipping introduced ties among 80 pixels. Pixel-level correlations were treated as descriptive because neighboring pixels were spatially correlated.

### Statistical analysis

The participant was the biological unit for between-participant inference. Deposits were treated as paired units only for within-participant registered analyses, and fields and pixels were nested observations. Participant-, field- and region-level summaries are reported as medians and ranges unless otherwise specified in the figure legends. Paired region-level comparisons within each registered section used exact two-sided Wilcoxon signed-rank tests (n = 6 pairs per section). Because the regions in each section were derived from a single participant, these tests were interpreted as measures of within-section consistency rather than participant-level inference. Severe and amyloid- negative participant summaries were compared using an exact two-sided Mann–Whitney U test (n = 3 and n = 5); the participant with mild deposition was excluded. Spearman coefficients and pixel-level AUCs were descriptive. Secondary descriptor, window and SNR analyses were exploratory; no multiplicity-adjusted inferential claims were made. The alpha level was 0.05.

## Data availability

De-identified data supporting the findings of this study are available from the corresponding author upon reasonable request, subject to institutional approval and restrictions protecting participant confidentiality.

## Code availability

Custom code used in this study is available from the corresponding author upon reasonable request.

## Acknowledgements

This work was supported by the Japan Society for the Promotion of Science (JSPS) KAKENHI (Grant Nos. 18H02000 and 24K03251, Grant-in-Aid for Scientific Research (B), and 21H04961, Grant-in- Aid for Scientific Research (A)), the Japan Science and Technology Agency (JST) Mirai Program (JPMJMI22G5), and the Japan Agency for Medical Research and Development (25ak0101251h0001) to H.K.; JSPS KAKENHI (Grant No. JP26KJ0047) to M.M.; the JSPS International Joint Research Program (JRP-LEAD with DFG; Grant No. 20261610) to R.S.; the JST Support for Pioneering Research Initiated by the Next Generation (SPRING; JPMJSP2123) to T.-H.C.; and the French National Research Agency through the Investments for the Future programme (ANR-10-LABX-0074, Sigma-LIM) and the French CNRS through International Research Project DEEP’nCARS to P.L. The authors thank J. Ukon of Ukon Craft Science, Ltd. for establishing the collaboration between the Japanese and French laboratories.

## Author contributions

T. Y. and H.K. conceived the study. K.O. and T.-H.C. performed the experiments. Y.M., M.M. and P.L. contributed to the development of the experimental system. T.Y., T.O. and K.N. provided human tissue and clinical or pathological annotation. K.O. developed the analysis. K.O., R.S. and H.K. interpreted the data. K.O. and H.K. drafted the manuscript. All authors critically revised and approved the final manuscript and agreed to be accountable for the work.

## Additional information

Supplementary Information is available for this paper. Correspondence and requests for materials should be addressed to H.K.

