## Supplemental Document and Figures for "Amide I kurtosis maps cardiac amyloid in human tissue by ultrabroadband multiplex coherent anti-Stokes Raman scattering (CARS) microscopy"

#### Registration diagnostics

For the primary CARS–histology section pair, the Congo red image was registered to the CARS field using 54 corresponding landmarks and a thin-plate-spline transform. The median displacement attributable to the non-affine component was 6.0 μm. The landmark convex hull covered 99.0% of the tissue mask and all registered deposit area. These values describe deformation and landmark coverage rather than registration error.

#### Replication in the second registered section

A registered CARS–histology section pair from a second participant, who was also included in the nine-participant analysis in Fig. 3, showed spatial correspondence between amide I kurtosis and Congo red-positive tissue (Spearman's ρ = 0.39; AUC = 0.73). In the discovery participant, apparent amide I FWHM was smaller in deposit-associated tissue than in paired surrounding myocardium in all six registered region pairs (exact two-sided Wilcoxon signed-rank test, P = 0.031; Supplementary Fig. S1). In the second participant, five of the six registered region pairs also showed a smaller apparent FWHM in deposit-associated tissue (median paired decrease, 17.8 $cm^{-1}$; P = 0.0625). Although registered histology was available for only two participants, the second pair showed similar spatial correspondence and deposit-associated band narrowing in a participant distinct from the discovery case.

### Window and SNR sensitivity

The kurtosis contrast persisted for windows centered at 1660 $cm^{-1}$ with half-widths of 40–100 $cm^{-1}$, but not at ±30 $cm^{-1}$, where the baseline endpoints intersected the band flanks (Supplementary Fig. S2). The primary analysis used a half-width of 60 $cm^{-1}$. Native field-level SNR ranged from 2.54 to 22.8 across fields acquired in two sessions. To assess sensitivity to signal quality, each field was numerically degraded to the lowest native SNR observed in the additional cohort (2.54). The gap between the severe and amyloid-negative kurtosis ranges decreased from 1.39 to 0.09 (Supplementary Fig. S3). The participant with mild deposition contributed three fields with kurtosis values of 1.65–2.47; one value fell below every amyloid-negative field (Fig. 3a). These data were not used to infer three-grade classification.

### MCR-ALS transfer and targeted spectral maps

The four-component MCR-ALS model converged in 28 iterations with 18.4% lack of fit ($R^2$ = 96.6%). In the second registered section, its component maps showed weaker correspondence with Congo red-positive tissue than the kurtosis map (spatial AUCs, 0.49–0.58 versus 0.73). Because MCR-ALS and kurtosis were derived from the same CARS data, their spatial correspondence provides internal consistency rather than independent molecular validation.

Guided by the deposit-associated component, we mapped the phenylalanine-associated band near 1003 $cm^{-1}$, the tryptophan-associated region near 1550 $cm^{-1}$ and the tyrosine doublet near 825 and 855 $cm^{-1}$ (Supplementary Fig. S4). Among the targeted aromatic indices, the phenylalanine/CH bending ratio showed the strongest correspondence with registered Congo red-positive tissue (AUC = 0.77; ρ = 0.52), followed by the tryptophan-associated 1550 $cm^{-1}$-region/CH bending ratio (AUC = 0.70; ρ = 0.38).

The tyrosine-band index showed an inverse association (AUC = 0.31; ρ = −0.33). These maps were derived from the same CARS dataset as the kurtosis map and provide complementary chemical context.

### Supplementary figure legends

Supplementary Figure S1. Apparent amide I FWHM in two CARS–histology section pairs from two participants. Pseudo-Voigt FWHM was obtained from the mean spectrum of each deposit-associated CARS region and its paired 20–60-pixel surrounding-myocardium annulus; lines connect paired regions. Regions were defined by Otsu thresholding of the registered Congo red image (connected regions ≥ 400 pixels). In the discovery participant (first pair, n = 6 regions), FWHM was lower in all six deposit-associated regions (medians, 34.7 versus 48.5 cm$^{-1}$; median paired decrease, 13.8 cm$^{-1}$; exact two-sided Wilcoxon signed-rank test, P = 0.031). In the second participant (second pair, n = 6), FWHM was lower in five of six regions (medians, 28.8 versus 47.0 cm$^{-1}$; median paired decrease, 17.8 cm$^{-1}$; P = 0.0625). The exception had the smallest surrounding-tissue annulus (272 pixels).

Supplementary Figure S2. Dependence of amide I kurtosis on analysis-window width. Field-median kurtosis values for severe, mild and amyloid-negative tissue were recalculated using windows centered at 1660 cm$^{-1}$ with half-widths of 30–100 cm$^{-1}$. The primary ±60 cm$^{-1}$ window is indicated. Separation between severe and amyloid-negative fields was retained at half-widths of 40–100 cm$^{-1}$ but not at ±30 cm$^{-1}$, where the window endpoints intersected the amide I band flanks. Fifteen fields from nine participants are shown.

Supplementary Figure S3. Sensitivity of amide I kurtosis to numerical SNR degradation. Field-median kurtosis values for severe and amyloid-negative tissue are shown before and after numerical degradation to an SNR of 2.54, the lowest native field-level SNR among the 15 additional-cohort fields. The gap between the group ranges decreased from 1.39 to 0.09. This exploratory analysis was not used to select a threshold or estimate diagnostic performance.

Supplementary Figure S4. Targeted aromatic-region maps in the primary registered field. (a) Phenylalanine-associated band area over 996–1010 cm$^{-1}$ divided by the CH bending-band area near 1450 cm$^{-1}$. (b) Tryptophan-associated band area over 1540–1565 cm$^{-1}$ divided by the same CH bending-band area. (c) Tyrosine-band index, $I_{855}/(I_{825} + I_{855})$, calculated using the 848–862 and 818–832 cm$^{-1}$ windows. Band areas in (a) and (b) were integrated after local linear baseline correction. (d) Descriptive spatial correspondence with the registered Congo red-positive mask: amide I kurtosis, AUC = 0.79 and ρ = 0.53; phenylalanine/CH bending, AUC = 0.77 and ρ = 0.52; tryptophan-associated region/CH bending, AUC = 0.70 and ρ = 0.38; tyrosine-band index, AUC = 0.31 and ρ = −0.33. All values are in-sample and exploratory (n = 54,815 spatially correlated pixels in one field). Scale bars, 200 µm.

Supplementary Figure S1

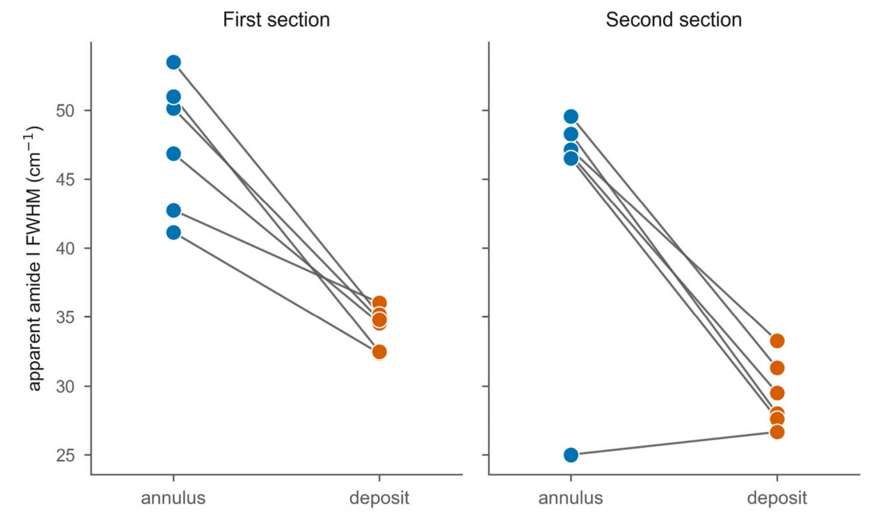


Supplementary Figure S2

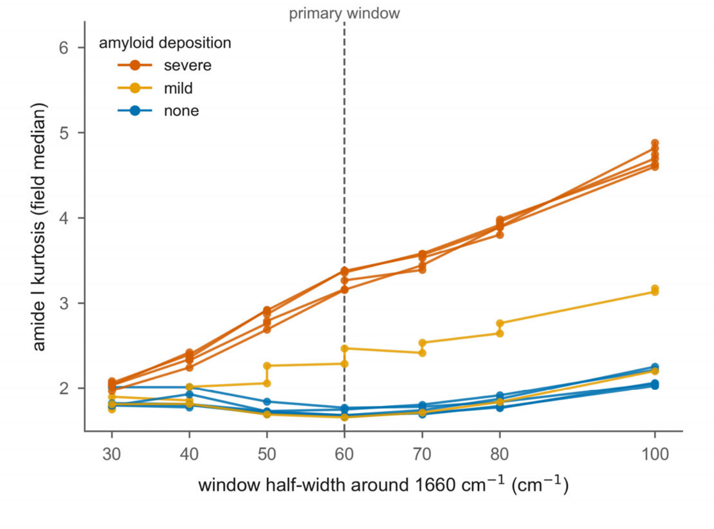

Supplementary Figure S3

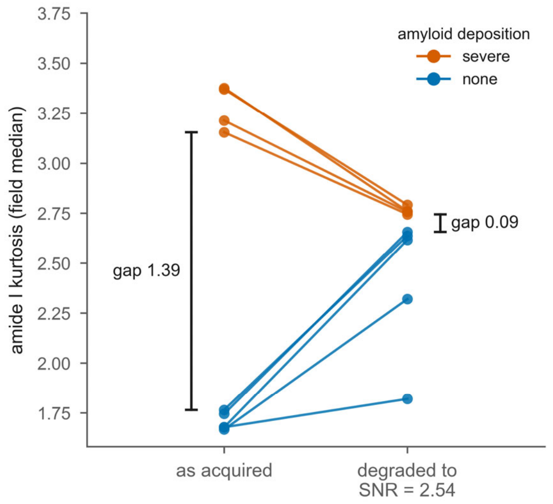


Supplementary Figure S4

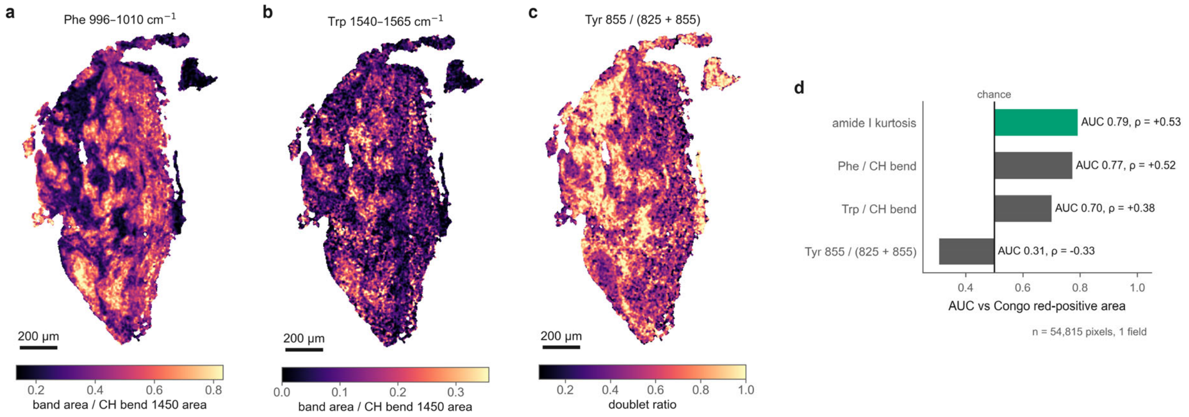
